## Supplementary information for "Brown Adipose Tissue and Skeletal Muscle Coordinately Contribute to Thermogenesis in Mice"

Yuna Izumi-Mishima, Rie Tsutsumi, Tetsuya Shiuchi, Saori Fujimoto,

Momoka Taniguchi, Mizuki Sugiuchi, Manaka Tsutsumi,

Yuko Okamatsu-Ogura, Takeshi Yoneshiro,Masashi Kuroda,

Kazuhiro Nomura, and Hiroshi Sakaue

**Supplementary Table 1. Proportion of immobilized skeletal muscle weight for cast-immobilized mice.** Body weight and tissue weights for skeletal muscle of the posterior cervical region of 12-week-old male C57BL/6J mice are shown.

| Mouse | Body weight (g) | Hind limbs (g) | Fore limbs (g) | Other (g) | Total (g) | Hind limbs (%) |
| --- | --- | --- | --- | --- | --- | --- |
| 1 | 27.93 | 1.762 | 0.454 | 1.256 | 3.472 | 50.74885 |
| 2 | 28.58 | 1.516 | 0.517 | 1.560 | 3.593 | 42.19315 |
| 3 | 30.06 | 1.958 | 0.476 | 2.256 | 4.690 | 41.74840 |
| 4 | 31.66 | 1.805 | 0.536 | 1.797 | 4.138 | 43.62011 |
| Average | 29.5575 | 1.76025 | 0.49575 | 1.71725 | 3.97325 | 44.57763 |

**Supplementary Table 2. Soleus and gastrocnemius tissue weights for individual mice subjected to iBAT removal (or sham surgery) followed by cast immobilization for up to 7 days.**

| Sham (mg) | | | | Removed (mg) | | | |
| --- | --- | --- | --- | --- | --- | --- | --- |
| Group | No. | Soleus | Gastrocnemius | Group | No. | Soleus | Gastrocnemius |
| Control | 1 | 23 | 333.4 | Control | 1 | 17.6 | 310.7 |
|  | 2 | 20.4 | 299.1 |  | 2 | 17.4 | 271.1 |
|  | 3 | 18.2 | 303.4 |  | 3 | 16.1 | 229.5 |
|  | 4 | 17.9 | 308.4 |  | 4 | 18.4 | 287.8 |
|  | 5 | 18.9 | 302 |  | 5 | 20.2 | 283.9 |
|  | 6 | 17.6 | 289.5 |  | 6 | 14.5 | 304.5 |
|  | 7 | 21.6 | 285.9 |  | 7 | 16.7 | 281.8 |
| 10H | 1 | 22.7 | 316.3 | 10H | 1 | 23.9 | 320 |
|  | 2 | 17.9 | 284.8 |  | 2 | 18.9 | 309 |
|  | 3 | 16.6 | 268.2 |  | 3 | 15.9 | 270.7 |
|  | 4 | 15.3 | 253.6 |  | 4 | 16.3 | 268.8 |
|  | 5 | 18.3 | 286.8 |  | 5 | 16.8 | 278.5 |
| 24H | 1 | 21.2 | 296.1 | 24H | 1 | 18.2 | 348.3 |
|  | 2 | 18.5 | 296.5 |  | 2 | 13.1 | 277 |
|  | 3 | 18.5 | 278.1 |  | 3 | 15.5 | 279.4 |
|  | 4 | 18 | 273 |  | 4 | 16.6 | 276.9 |
|  | 5 | 15.2 | 271.4 |  | 5 | 16.5 | 272.8 |
| Day3 | 1 | ND | 321.8 | Day3 | 1 | 18.8 | 372.4 |
|  | 2 | 20.1 | 271.6 |  | 2 | 19.4 | 301 |
|  | 3 | 14 | 277.7 |  | 3 | 19.1 | 291.1 |
|  | 4 | 15.8 | 253.3 |  | 4 | 15.2 | 254.3 |
| Day5 | 1 | 17.4 | 278.4 |  | 5 | 17.6 | 275.5 |
|  | 2 | 14.1 | 259.5 | Day5 | 1 | 16.2 | 286.4 |
|  | 3 | 14.8 | 273.7 |  | 2 | 16.7 | 238.3 |
|  | 4 | 14 | 200.8 |  | 3 | 14.9 | 278.5 |
|  | 5 | 10.7 | 213.3 |  | 4 | 16.5 | 222.2 |
| Day7 | 1 | 14.4 | 256.6 |  | 5 | 13.4 | 262.8 |
|  | 2 | 15.4 | 252.7 | Day7 | 1 | 19 | 287.8 |
|  | 3 | 14.4 | 239.8 |  | 2 | 18 | 251.4 |
|  | 4 | 14.8 | 261.3 |  | 3 | 16.7 | 235.8 |
|  | 5 | 16 | 268.2 |  | 4 | 11 | 239.2 |
|  | 6 | 15.6 | 264.5 |  | 5 | 12.7 | 234.4 |
|  |  |  |  |  | 6 | 11.8 | 244.8 |

**Supplementary Table 3. Sequences (5′→3′) of PCR primers.**

| Transcript  (Mouse) | Forward primer | Reverse primer |
| --- | --- | --- |
| *Gapdh* | AAAATGGTGAAGGTCGGTGTG | TTGACTGTGCCGTTGAATTTG |
| *Ucp1* | CTCAGCCGGAGTTTCAGCTT | GTTTTTGCCAGGGTGGTGAT |
| *Ucp2* | TCTTGCCGATTGAAGGTCCC | CTAGCCCTTGACTCTCCCCT |
| *Ucp3* | TGTCTCTGCCTTTGGAGCTG | GGCCCTCTTCAGTTGCTCAT |
| *Sln* | AGGGGCCATGCTATACTCCA | TGGGCAGCCTACAAGAACAG |
| *Camk2a* | GGTCAGGAGTATGCTGCCAAG | CCCACCAGTAACCAGATCGAA |
| *Fbxo32* | AGGAGCGCCATGGATACTGT | GAAGTTCTTTTGGGCGATGC |
| *Trim63* | GACAGTCGCATTTCAAAGCA | AACGACCTCCAGACATGGAC |
| *Ppargc1a* | TCACACCAAACCCACAGAAA | TCTGGGGTCAGAGGAAGAGA |
| *Tfam* | CGGCTCAGGGAAAATTGAAG | AGCCATCTGCTCTTCCCAAG |
| *Bcat2* | TTCCAGAACCTCACGCTACAC | TAGCAGAACGTAGCATCCTGTC |
| *Bckdha* | AGGAGGTGCTGAAGTTCTACC | CGCCATAGTTGGTCATGTAGAAG |
| *Slc1a5* | GCAGTGCACCAACCAAAGAG | CCAGGCCCAGGATGTTCATT |
| *Slc7a5* | TTTTGCTCGGCTTCATCCAG | ACAACTTCTGCTGCAGGTTG |
| *Slc38a2* | CCTTCTGGTGTCCCTTGTCC | CTGCGGTGCTATTGAATGCC |
| *Slc43a1* | CCTGGGCCTCCTACTTCTCT | TGCAGGTAGAAAGCCACAGG |
| *Slc25a10* | CAGGATGCAGAACGACATGAA | ACCATCCAGGGCATGAGAGTA |
| *Slc25a44* | TCGCTGCTAACGTACATCCC | AGACAATGTGAGGGCACTCC |
| *Cd36* | GCCAAGCTATTGCGACATGA | AAAGGCATTGGCTGGAAGAA |
| *G6pc1* | AAGACTCCCAGGACTGGTTCATCC | TAGCAGGTAGAATCCAAGCGCG |
| *Pck1* | TGCTGATCCTGGGCATAACTAACC | TGGGTACTCCTTCTGGAGATTCCC |
| *Il1b* | TGGCAACTGTTCCTG | GGAAGCAGCCCTTCATCTTT |
| *Il6* | CACAGAAGGAGTGGCTAAGGACCA | ACGCACTAGGTTTGCCGAGTAGA |
| *Il15* | ACATGGCCCTCTGGCTCTT | AGCTGCCATCCATCCAGAA |
| *Tnfa* | TGAACTTCGGGGTGATCGGT | GTTTGCTACGACGTGGGCTAC |
| *Mcp-1* | CTGTTCACAGTTGCCGGCTG | AGCTTCTTTGGGACACCTGCT |
| *Irisin* | GAAGGAGATGGGGAGGAACC | GGTGTGCTGGTTTCTGATGC |
| *Fgf21* | GGATCGCCTCACTTTGATCC | ATCCTGGTTTGGGGAGTCCT |
| *Saa3* | ATGCTCGGGGGAACTATGAT | TCCATGTCCCGTGAACTTCT |
| *Socs3* | CTTTTCTTTGCCACCCACGG | CGACAAAGATGCTGGAGGGT |
| *Crh* | TCAGAGCCCAAGTACGTT | AGGGACTTCTCTCAGGAT |
| *Bmp8b* | CCTCGAACAGCAAGACCACT | GCACTCCCCAAGCACAGTAAT |
